# Perchlorate-Specific Proteomic Stress Responses of *Debaryomyces hansenii* Could Enable Microbial Survival in Martian Brines

**DOI:** 10.1101/2022.05.02.490276

**Authors:** Jacob Heinz, Joerg Doellinger, Deborah Maus, Andy Schneider, Peter Lasch, Hans-Peter Grossart, Dirk Schulze-Makuch

## Abstract

If life exists on Mars, it would face several challenges including the presence of perchlorates, which destabilize biomacromolecules by inducing chaotropic stress. However, little is known about perchlorate toxicity for microorganism on the cellular level. Here we present the first proteomic investigation on the perchlorate-specific stress responses of the halotolerant yeast *Debaryomyces hansenii* and compare these to generally known salt stress adaptations. We found that the responses to NaCl and NaClO_4_-induced stresses share many common metabolic features, e.g., signaling pathways, elevated energy metabolism, or osmolyte biosynthesis. However, several new perchlorate-specific stress responses could be identified, such as protein glycosylation and cell wall remodulations, presumably in order to stabilize protein structures and the cell envelope. These stress responses would also be relevant for life on Mars, which - given the environmental conditions - likely developed chaotropic defense strategies such as stabilized confirmations of biomacromolecules and the formation of cell clusters.

## Introduction

Life as we know it requires energy as well as access to CHNOPS (carbon, hydrogen, nitrogen, oxygen, phosphorus, sulfur), trace elements, and liquid water. On Mars, energy would be provided to putative life chemically or via sunlight, carbon is accessible through the thin but CO_2_-rich atmosphere, and other essential elements are abundant in the regolith (1). Availability of liquid water, however, is strongly restricted due to the low atmospheric pressure of approx. 6 mbar and mostly subzero temperatures on Mars (2). One of the few possibilities to generate liquid water in the Martian near surface is the formation of temporarily stable brines via deliquescence, a process in which a hygroscopic salt absorbs water from the atmosphere and dissolves within that water (2). It has been shown that deliquescent water is sufficient to drive the metabolism of halotolerant methanogenic archaea (3). Intriguingly, several hygroscopic salts have been detected on Mars (4). Among those are very deliquescent and freezing point depressing perchlorates (CLO_4_^-^), which are widely distributed on the Martian surface (5) but appear in natural environments on Earth only occasionally in hyperarid deserts (6, 7).

Brines formed via deliquescence provide diverse challenges for microbial life. High salt concentrations lead to osmotic stress and reduce water activity, which is a measure for the amount of unbound water molecules in a solution available for biological processes (8). Furthermore, salts can induce ion-specific stresses like interferences with the cell’s metabolism or changes in cell permeability through variations in ionic hydration shells (9). Some anions like perchlorate additionally evoke chaotropic stress (10), i.e., they destabilize biomacromolecules like proteins, presumably through nonlocalized attractive dispersion forces (11). In *Pseudomonas putida*, it has been shown that chaotropic solute-induced water stress mainly leads to upregulation of proteins involved in stabilization of biological macromolecules and membrane structure (12). However, detailed research on microbial responses to perchlorate-induced chaotropic stress is still lacking.

Here, we present a proteomic study investigating the perchlorate-specific stress response on *Debaryomyces hansenii* to evaluate the physiological adaptations required for microorganisms to thrive in the Martian near surface. The halotolerant yeast *D. hansenii* has been chosen as a model organism as it has been described earlier to tolerate the highest perchlorate concentrations reported to date (13, 14). This yeast provides a large metabolic toolset to counteract salt stress, such as the high-osmolarity glycerol (HOG) pathway which enables stress signaling and concomitant biosynthesis of the osmoprotectant glycerol (15). Its close relation to the intensively studied bakery yeast *Saccharomyces cerevisiae* greatly facilitates the annotation of proteins and thus prediction of their functions. For the investigation of the proteome of *D. hansenii*, we choose a recently developed proteomics protocol called SPEED (Sample Preparation by Easy Extraction and Digestion) which enables sample-type independent deep proteome profiling with high quantitative accuracy and precision (16, 17).

This is the first study investigating perchlorate-specific stress responses (i.e., with a significant distinction compared to general salt stress) with an untargeted proteomic approach to provide novel and fundamental understanding of the required cellular adaptation mechanisms for life in perchlorate-rich, chaotropic habitats on Earth, Mars, and beyond.

## Results

In order to distinguish the perchlorate-specific stress response of *D. hansenii* from general osmotic and salt stress responses proteomes of cell cultures containing either NaClO_4_, NaCl or no additional salts in growth medium DSMZ #90 were analyzed. Two different stress regimes were investigated (Fig. 1). At moderate salt stress (1.5 m NaClO_4_ and 2.4 m NaCl, with m = molality [mol/kg]), growth media were inoculated with a salt-free culture to provoke a salt shock response. Cell growth at high salt stress conditions (2.4 m NaClO_4_ and 3.9 m NaCl) could only be enabled by long-term adaption of cells to stepwise increasing salt concentrations (Fig.1). All samples were prepared as biological triplicates and cells were harvested in the late exponential growth phase (one day for salt-free treatment, three days for 1.5 m NaClO_4_ and 2.4 NaCl, six days for 2.4 m NaClO_4_, and seven days for 3.9 m NaCl, coinciding with a much slower growth at increasing salt stress levels).

**Figure 1.**
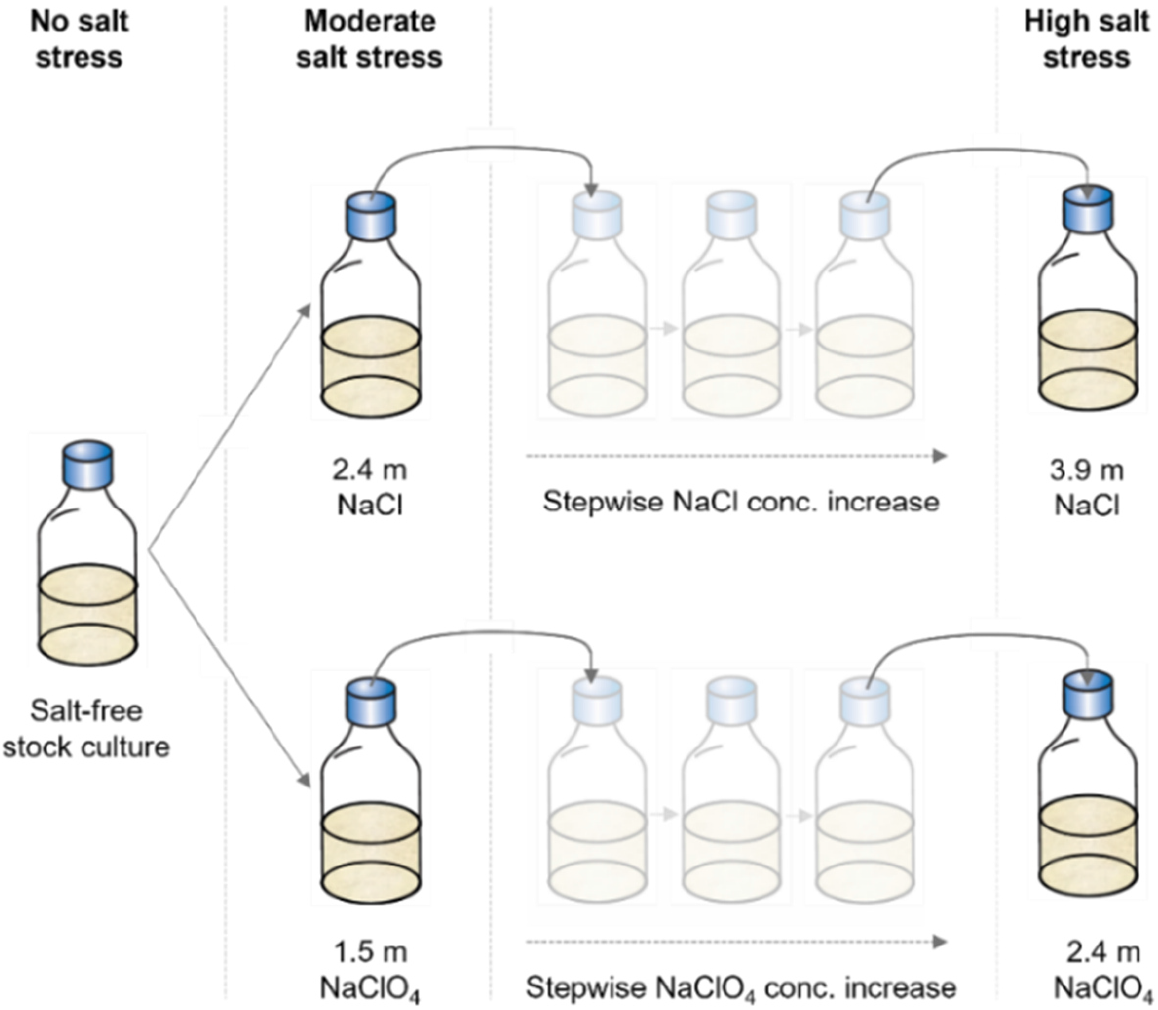
Workflow of the inoculation procedure and culture preconditioning. A salt-free stock culture of *D. hansenii* was frequently reinoculated into fresh growth medium. An aliquot of this culture was used to inoculate growth media exhibiting moderate cellular salt stress (2.4 m NaCl and 1.5 m NaClO_4_). To obtain cell growth at even higher salt concentrations, a stepwise concentration increase was needed for each inoculation step. The maximum salt concentrations used in this study were 3.9 m NaCl and 2.4 m NaClO_4_. Samples for protein extraction were taken in the late exponential growth phase of the respective treatments. Each treatment type was inoculated and treated in biological triplicates.

In total, 2713 proteins were detected representing a bulk coding sequence coverage of approx. 43*%*. Through analysis of variance (ANOVA, FDR < 0.01) of the z-normalized protein abundances, the expression of 1099 proteins was found to be significantly different between the five different treatment types (one salt-free control, and four salt-containing treatments). The salt stress level (moderate vs. high salt stress) had a stronger impact on the intensity of protein expression than the type of anion or molal concentration (2.4 m NaCl vs. 2.4 m NaClO_4_) as can be seen from the comparison of protein abundances of all replicates, which show similar protein expressions for the same salt stress regimes (Fig. 2a). This is confirmed by the principal component analysis (PCA) which revealed a clear clustering of the replicates of each treatment in dependence on salt stress level and type of anions (Fig. 2b). While the physiological response to different salt stress levels clustered along principal component 1 and explains 55% of the observed differences, the salt species had a lower impact on the variability (15%), as treatments exposed to chloride or perchlorate spread along the principal component 2.

**Figure 2.**
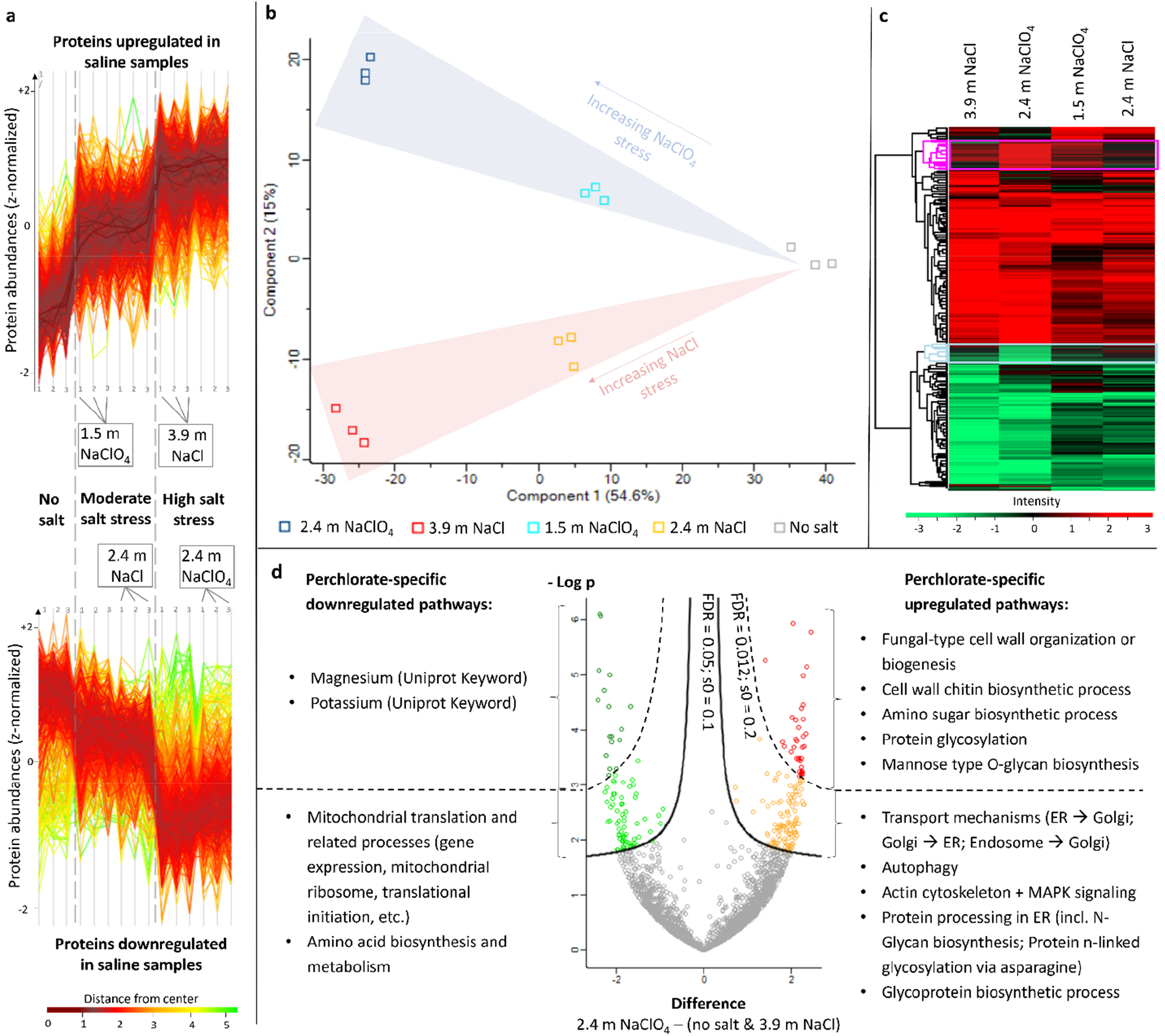
Results of the proteomic analyses. **(a)** Abundances of upregulated (upper plot) and downregulated (lower plot) proteins expressed in all investigated samples (three replicates for each treatment). **(b)** Principal component analysis (PCA) demonstrating clear clustering of all biological triplicates in dependence of salt stress level and type of anion. **(c)** Heat map including all proteins passing ANOVA (FDR < 0.01) and post hoc test (FDR < 0.05) generated by the Perseus software after hierarchical clustering. Upregulated proteins (compared to the salt-free treatment) are colored red and downregulated proteins are shown in green. Two exemplarily perchlorate-specific clusters are highlighted in pink for upregulated and in cyan for downregulated proteins. **(d)** Volcano plot visualizing perchlorate-specific regulated proteins with a high (FDR ≤ 0.012) and a lower significance (0.012 ≤ FDR ≤ 0.05). Significantly regulated metabolic pathways are analyzed with the STRING database.

Post hoc testing (FDR < 0.05) revealed 1068 proteins to be significantly regulated in at least one of the salt-containing samples compared to the salt-free treatment. The log2-fold changes of proteins in the saline treatments compared to the salt-free control were plotted in a heatmap with upregulated proteins colored red and downregulated protein shown in green (Fig. 2c). The heatmap reveals two major clusters, one containing the proteins predominantly upregulated in all saline treatments compared to the salt-free control, and a second cluster with downregulated proteins. This indicates that the overall salt stress response is relatively similar in all treatments. However, both main clusters also contain proteins that are substantially more regulated in the 2.4 m NaClO_4_ than in all other treatments. The two largest subclusters containing proteins of this category are highlighted in pink (for upregulated proteins) and cyan (for downregulated proteins) in Fig. 2c. Proteins in these subclusters represent the perchlorate-specific stress response, which apparently manifests only at high perchlorate concentrations, as protein expression patterns in the 1.5 m NaClO_4_ treatment are coinciding more with the NaCl than with the 2.4 m NaClO_4_ treatment.

This enables the possibility to investigate the significance of perchlorate-specific protein expression patterns by a volcano plot that presents the differences of the z-normalized protein abundances in the 2.4 m NaClO_4_ treatment and the control samples (salt-free and 3.9 m NaCl) vs. the logarithmic p value after a t-test (Fig. 2d). The resulting perchlorate-specific regulated proteins include the ones from the heatmap subclusters (marked pink and cyan in Fig. 2c) and additional proteins from smaller subclusters. Proteins were fed into the STRING database (18) which predicts physical and functional protein-protein interactions and identifies significantly enriched (FDR < 0.05) metabolic pathways that assort into protein clusters (Fig. 3a, and Table S1). The physiological interpretation of the perchlorate-specific enriched pathways is summarized in Fig. 3b and discussed in detail in section 3.2.

**Figure 3.**
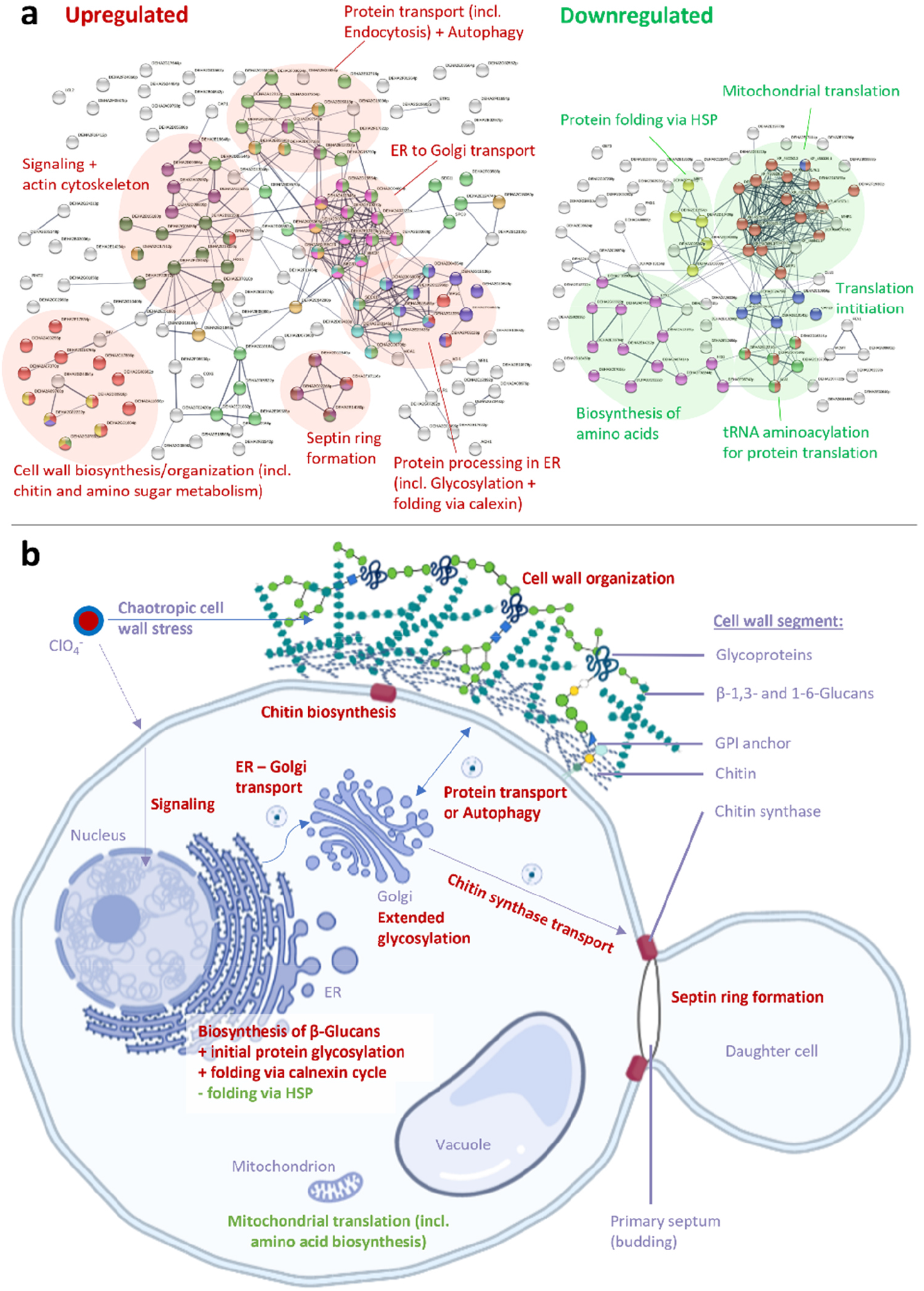
Perchlorate-specific stress responses. **(a)** STRING database calculated interactions of all upregulated (left) and downregulated (right) proteins involved in the perchlorate-specific stress response according to the volcano plot (Fig. 2d). The line thickness of the network edges indicates the strength of data support. The minimum required interaction score was set to 0.6 for up- and 0.5 for downregulated proteins. Colored proteins indicate significantly enriched metabolic pathways (FDR < 0.05) and are annotated in Table S1. The most prominent pathways are encircled. **(b)** A mother cell and a budding daughter cell of *Debaryomyces hansenii* displaying the most relevant metabolic pathways with perchlorate-specific upregulations (red) and downregulations (green) as explained in the main text. Created with BioRender.com.

In order to better evaluate the specificity of perchlorate-induced stress responses, they must be compared to the overall salt stress response which is shared by all saline treatments. For this purpose, proteins that show significant expression (FDR < 0.05) in all salt-containing treatments compared to the salt-free control have been analyzed with the STRING database. The results of this approach are discussed below and summarized in the Table S1 and Fig. S1.

## Discussion

### 3.1 General salt stress response

The general salt stress response observed in all saline treatments encompasses several metabolic stress response pathways, previously well described for *D. hansenii* and its close relative, the intensively investigated yeast *S. cerevisiae (15, 19)*. Only the most prominent pathways detected in this study are described below. Any form of environmental stress in yeasts is usually communicated from the cell envelope to the nucleus via signaling pathways such as mitogen-activated protein kinase (MAPK) pathways (20) accompanied by a rearrangement of cytoskeletal arrays (21). Several proteins involved in these signaling pathways were found to be upregulated in all saline treatments (see Table S1 and Fig. S1). For example, the upregulated serine/threonine-protein kinase CLA4 (DEHA2B12430p) modulates the expression of biosynthesis of glycerol (22), an important osmoprotectant in *D. hansenii* (15).

As a consequence of the received signal, the cell’s energy metabolism is upregulated to induce certain stress responses. In our experiments, several energy-releasing pathways were upregulated in the general osmotic stress response, such as the TCA cycle. Furthermore, processes in the peroxisomes showed upregulation including the energy-releasing beta-oxidation of fatty acids indicated by upregulation of the multifunctional beta-oxidation protein DEHA2A08646p, the acyl-coenzyme A oxidase POX1 (DEHA2D17248p), and the peroxisomal long-chain fatty acid import protein DEHA2B08646p. In yeasts, also the glyoxylate cycle, a variation of the TCA cycle (23), takes place in the peroxisomes as well as in the cytoplasm, and correspondingly the two key-enzymes, i.e., malate synthase (DEHA2E13530p) and isocitrate lyase (ICL1 DEHA2D12936p), are upregulated.

The energy released from these processes is presumably needed to guarantee survival under enhanced osmotic cell stress. For example, the amount of biosynthesized glycerol is increased as can be observed by the upregulation of the glycerol lipid metabolism which includes proteins that participate in the formation of glycerol, e.g. several significantly enriched glycerol-3-phosphate dehydrogenase complex proteins. In addition to the accumulation of glycerol, osmotic stress in *D. hansenii* can also be antagonized through ion transmembrane transporters (24). Even though not being incorporated in a significant enrichment, we found several ion transporter proteins to be upregulated, such as the ATPase-coupled cation transmembrane transporters DEHA2G09108p and DEHA2C02552p (Table S1). These two cation transporters showed the highest significance upon all proteins upregulated in the saline treatments (see volcano plot in Fig. S1). The provision of sufficient ATP required for the functioning of these transporters constitutes to the enhanced cellular energy demand.

Osmotic stress usually induces oxidative stress to a some extent, e.g. by production of reactive oxygen species (ROS) in the mitochondria (25). Hence, it is expected that proteins involved in oxidative stress responses are regulated under salt stress conditions as well. Indeed, we found antioxidant activity and glutathione metabolic processes to be upregulated, including the enzymes catalase (DEHA2F10582g) and peroxidase (DEHA2A02310p) enabling cell protection against oxidative stress.

The significantly enriched pathway forming the most pronounced and condensed upregulated protein cluster contains proteins involved in the modification-dependent protein catabolic process and ubiquitin mediated proteolysis (Fig. S1). Osmotic and induced oxidative stresses can cause protein misfolding (26). Proteins which cannot be refolded by chaperones are degraded by the proteosome of the cell (27). The so generated amino acids can then be reused for cellular amino acid metabolism, which forms a protein subcluster interwoven with the TCA cycle, indicating that the amino acids liberated by proteolysis feed the energy metabolism. Additionally, the recycling of amino acids via proteolysis conserves energy as compared to amino acid *de novo* biosynthesis.

Most of the stress responses described above require protein transport, e.g., for posttranslational modifications in the ER or the Golgi apparatus, for the transfer of proteins to their place of activity, or for the excretion of cell wall proteins. Consistently, many of the upregulated proteins in the general salt response are involved in protein transport mechanisms.

More than half of the proteins downregulated due to general salt stress are structural ribosomal constituents and have translation factor activity or are involved in the cytosolic (pre)ribosome biogenesis and related pathways such as the nucleotide metabolism (Fig. S1). Ribosome biogenesis is a complex and very energy-demanding process (28). Consequently, for saving energy, ribosome biogenesis is downregulated under various stress conditions (29). A transient reduction in ribosome biogenesis and translation together with the accumulation of glycerol has also been detected in *Candida albicans* upon salt stress (30). While ribosome biogenesis was generally downregulated in our experiments, we consistently observed upregulation of the ribosome-recycling factor RRF1 (DEHA2F14630g) which allows the ribosome to unbind from mRNA after the release of the generated polypeptide and to be reused for new translation processes instead of the energy-consuming *de novo* biosynthesis of ribosomes (31).

Upregulation of ribosome synthesis occurs only in response to favorable growth conditions and enables the cell to grow faster (32), while downregulation of translation via depletion of the ribosomal population is known to prolong the lifespan of cells (33). Consistently, we found that the downregulation of ribosome biosynthesis and concomitant translational processes coincided with slower cell growth under salt stress conditions.

The only other significantly enriched downregulated pathway forming a protein cluster that is physiologically not directly connected to the ribosome assembly and translational processes is the biosynthesis of ergosterol (Fig. S1), a component of fungal cell membranes (34). In *S. cerevisiae*, the downregulation of the ergosterol biosynthesis has already been described earlier as response to hyperosmotic stress (35). It has been hypothesized that it results from increased uptake of Na^+^ and/or a decreased Na^+^ extrusion in a plasma membrane environment with elevated levels of ergosterol (35). Furthermore, sterol biosynthesis is a highly energy-consuming process (36), and its downregulation might constitute, similar to the downregulation of the ribosome biogenesis, an energy-saving approach.

### 3.2 Perchlorate-specific stress response

In a previous study we demonstrated that *D. hansenii* has the highest microbial perchlorate tolerance reported to date (14). However, subsequent experiments revealed that the tolerance towards NaClO_4_ (2.5 mol/kg) was still more than one third lower than towards NaCl (4.0 mol/kg), even though the water activity was substantially higher in the NaClO_4_-containing growth medium (0.926) than in the NaCl-rich medium (0.854) (13). Differences in salt tolerances were interpreted by the authors to account for the chaotropic stress exerted by the perchlorate anion. This interpretation is strongly supported by our proteomic investigations as explained below.

As a chaotropic ion, perchlorate is destabilizing biomacromolecules such as proteins (37) or glycan (i.e. polysaccharide) macromolecules (38). The fungal cell wall of *D. hansenii*’s close relative, *S. cerevisiae*, consists of approx. 85% glycans (incl. 1-2% chitin) and 15% cell wall proteins (39). Hence, it can be expected that the presence of chaotropic perchlorate induces cell wall stress in addition to the general salt stress. Our results suggest that yeast cells counteract this chaotropic stress to a certain extent by increasing the bioproduction rate of cell wall components.

For example, chitin metabolic processes (incl. the synthesis of its amino sugar precursors) were found to be significantly upregulated under perchlorate stress conditions (Fig. 3) and show a higher significance for the perchlorate-specific stress response than other metabolic pathways (Fig. 2d). Although being a minor component of fungal cells wall, chitin provides important structural stability (40). Chitin is produced by chitin synthases, such as the upregulated DEHA2D03916p, to a lesser degree directly in the lateral cell wall and to a higher extent in the primary septum during cell budding (39), presumably to protect the emerging nascent cell (40). This highlights the importance of enriched chitin synthesis already during budding under perchlorate stress which otherwise might chaotropically destabilize the nascent cell envelope. This also explains the upregulation of septin proteins (Fig. 3) which provide structural support during cell division at the septum (41).

Another metabolic pathway upregulated under perchlorate-specific stress conditions is the glycosylation of proteins. Studies have shown that N-glycosylation is stabilizing proteins (42) also with respect to chaotropic denaturation (43). The glycosylation-induced increase in protein stability affects both intracellular proteins as well as cell wall proteins. However, in contrast to intracellular proteins which are usually N-glycosylated in the ER with only 9 to 13 glycan residues, cell wall proteins experience an extensive additional glycosylation (including O-glycosylation) in the Golgi apparatus resulting in a highly branched structure containing as many as 200 glycan residues (39). We found that one of the most pronounced and densest perchlorate-specific upregulated protein clusters contained proteins involved in the ER to Golgi vesicle mediated transport (Fig. 3a). This suggests that a large part of the upregulated glycosylation processes is applied to cell wall proteins. Furthermore, three of the upregulated proteins involved in protein glycosylation are O-mannosyltransferases involved in O-glycosylation, which is essential for cell wall rigidity (44) and also upregulated upon heat stress (45).

The glycosylated cell wall proteins are transported from the Golgi apparatus via vesicle mediated transport to the cell wall. A protein that needs to be highlighted in this context is the upregulated Chs5-Arf1p-binding protein DEHA2G07832p whose homologue in *S. cerevisiae* mediates export of chitin synthase 3 from the Golgi apparatus and the transport to the plasma membrane in the bud neck region (46) confirming the importance of cell wall chitin metabolic processes under perchlorate stress. Misfolded proteins or proteins chaotropically denatured despite stabilizing glycosylation might be autophagically degraded explaining the upregulation of proteins involved in autophagy (Fig. 3).

Among the perchlorate-specific upregulated proteins are several proteins that are involved in cell wall biogenesis and remodulation. Apart from the chitin synthases, these are, e.g., the two glycosidases DEHA2G21604p (glycoside hydrolase family 16, CRH1 homolog in *S. cerevisiae*) and DEHA2G18766p (Glucan 1,3-beta-glucosidase BGL2) responsible for glucan cross-linking and chain elongation in the cell wall (47, 48). Stronger cross-linking of cell wall components and concomitant disability to separate cells after cell division might explain the formation of cell chains of *Hydrogenothermus marinus* (49) and of large cell aggregates of *Planococcus halocryophilus* (50) when exposed to perchlorate stress.

In comparison to the upregulated cell wall biosynthesis and organization as well as the protein glycosylation, all downregulated perchlorate-specific processes have a lower significance (Fig. 2d). The most pronounced downregulated protein subcluster contains proteins involved in mitochondrial translation and is physiologically linked to the simultaneously downregulated amino acid biosynthesis, the tRNA aminoacylation for protein translation, and the translation initiation (Fig. 3a). Apart from energy-saving aspects similar to the downregulation of cytosolic translation under general salt stress conditions (section 3.1), it has been described that changes in mitochondrial translation accuracy modulate cytoplasmic protein quality control (51). For example, it has been observed that decreasing mitochondrial translation output coincides with cytoplasmic protein folding (52) which seems plausible under chaotropic stress conditions that promote destabilization of protein tertiary and quaternary structures.

Therefore, it might be surprising at first, that the machinery for protein folding via heat shock proteins (often acting as chaperons) is downregulated under perchlorate stress conditions (Fig. 3). However, protein folding is regulated by two major folding pathways. The general pathway is mostly mediated by 70-kDa heat shock proteins (Hsp70), while the second pathway, called the calnexin cycle, is dedicated for N-glycosylated proteins and requires, among others, the action of the proteins calnexin (or its homologue calreticulin) and disulfide isomerase (53). Since we observed a high degree of protein glycosylation in the perchlorate-specific stress response, it seems likely that not the heat shock protein mediated folding pathway is upregulated under perchlorate stress, but rather the calnexin cycle. Indeed, we detected perchlorate-specific upregulations of the disulfide isomerase DEHA2E23628p, and the calnexin homologue DEHA2E03146p as part of the of the protein processing in the ER cluster (Fig. 3a, Table S1).

The reduced mitochondrial translation might also be explained by the prevention of proteotoxic stress within the mitochondria as mitochondrial encoded proteins cannot be stabilized by glycosylation which takes place exclusively in the ER and Golgi apparatus (intramitochondrial glycosylation is under debate (54), however, this process has not yet been described for yeast cells), and therefore might be denatured more easily under perchlorate stress. Following this argumentation, the mitochondrial translation might be downregulated to the minimal required performance to avoid accumulation of denatured proteins within the mitochondria.

### 3.3 The role of perchlorate-induced oxidative stress

Due to the high oxidation state (+7) of the chlorine atom in the center of the perchlorate anion, it is expected that perchlorate exhibits a stronger oxidation stress response than observed in the general salt stress response. Indeed, there is evidence that several genes in microorganisms from sediments of hypersaline ponds increase both the resistance to perchlorate and to oxidative stress induced by hydrogen peroxide (55). Furthermore, increased levels of lipid peroxidation after growth of different species of cyanobacteria in perchlorate-containing growth media were interpreted as results of oxidative stress (56). However, the authors did not investigate whether the oxidative stress is perchlorate-specific or the result of general salt or osmotic stresses.

From all proteins potentially involved in oxidative stress response (e.g., superoxide dismutase, catalase, glutathione reductase, glutathione peroxidase, glutaredoxin, glyoxalase, or thioredoxin), in our experiments only the glutathione reductase GLR1 was observed to be perchlorate-specifically upregulated with a low significance (FDR > 0.012, Table S1). GLR1 is involved in a multiplicity of cellular functions including besides the protection of cells from oxidative damage also amino acid transport as well as DNA and protein synthesis (57). This indicates that the oxidative stress response is not substantially upregulated under perchlorate stress compared to general salt stress.

Reducing the mitochondrial translation activity (Section 3.2) might also be interpreted as an attempt of the cell to minimize ROS production during aerobic respiration. Indeed, previous studies indicated that perchlorate induces oxidative stress to mitochondria by enhanced ROS production (58, 59). Yet, similar to the enhanced lipid peroxidation of cells grown in perchlorate-containing medium (56), the missing comparisons with NaCl or other solutes made it impossible for the authors of these studies to proof that the increased ROS levels resulted from perchlorate-specific reactions and are not caused by the general osmotic stress. If the reduced mitochondrial translation activity observed in our experiments would be a result of an enhanced oxidative stress, a concomitant downregulation of respiratory chain proteins would be expected to occur. However, we did not observe a conclusive downregulation of these kind of proteins. For example, while the cytochrome c oxidase (COX) assembly mitochondrial protein DEHA2C13244p was downregulated, the COX subunit 9 was upregulated under perchlorate-specific stress conditions.

In summary, the proteomic data suggests that antioxidant activity is important for survival under general salt stress conditions (Section 3.1), but the oxidative stress induced specifically by perchlorate seems to play only a minor role compared to the chaotropic stress. This is in accordance with previous experiments demonstrating that the more oxidatively reactive (but less chaotropic) chlorate anion (ClO_3_^-^) can be better tolerated by *D. hansenii* than perchlorate, which indicates the oxidative character alone cannot account significantly to the additional stress exhibited by NaClO_4_ compared to NaCl (13). A possible explanation for this phenomenon is that perchlorate is astonishingly stable in solution under ambient temperatures (60) due to the reduction rate-limiting oxygen atom transfer (61). These additional stressors (chaotropicity, and potentially to a minor degree also oxidative stress) presumably require different or more distinct stress signaling which likely explains the upregulation of proteins involved in signaling and the actin cytoskeleton organization pathways (Fig. 3) in addition to the proteins expressed as general salt stress responses.

The most relevant of the above-described perchlorate-induced chaotropic stress responses are graphically summarized in Fig. 3b. In particular, cell wall genesis is a very energy-consuming process (62), but also intensive protein glycosylation required for protein stability under perchlorate stress costs additional energy compared to non-chaotropic conditions. While in the non-chaotropic NaCl-stressed samples most of the energy provided by stress-adapted cell metabolism can be used to counteract osmotic and induced oxidative stresses, in perchlorate-containing samples a substantial part of the cellular energy demand is required for counteracting chaotropic stress resulting in a lower NaClO_4_ tolerance of *D. hansenii* compared to NaCl.

### 3.4 Consequences for microbial habitability of perchlorate-rich environments on Mars

This study provides new insights for putative life on Mars if it exists in perchlorate-rich regions, which have been identified during the exploration of Mars (5, 63, 64). Protein glycosylation and cell wall organization are major stress responses emerging only after long-term adaptions to high perchlorate concentrations, while being not significantly expressed after perchlorate-shock at moderate salt concentrations. Hence, it is likely that biomacromolecules and cell envelops of putative Martian microorganisms exposed to perchlorate-rich brines would evolve stable confirmations and prefer covalent bounds and cross-linking over looser electrostatic interactions, hydrogen bonding or hydrophobic effects. Additionally, cell components susceptible to chaotropic stress might be stabilized by the attachment of polymers similar to stabilization effects via protein glycosylation as observed in our experiments.

Furthermore, previous microscopic observations (49, 50) indicate that larger cell aggregates are more likely to occur (possibly due to cell wall rearrangements and cross-linking) under perchlorate stress than single cells. Consequently, cell clusters or biofilms might be considered as potential macroscopic visible biosignatures on Mars, however, metabolomic changes under perchlorate stress should be investigated in upcoming experiments as well in order to identify potential perchlorate-specific biomarkers on the molecular level.

The presented results are also important for *in-situ* resource utilization (ISRU) technologies to support a human outpost on Mars (65). Oxygen and food production by phototrophic microorganisms and the recycling of waste material in perchlorate-rich Martian soil might be conducted by “chaotolerant” (66) organisms, because they would likely possess a metabolic toolset for stabilization of biomacromolecules similar to *D. hansenii*. Alternatively, genes responsible for increased biomacromolecular stability might be used in synthetic biology to create perchlorate resistance strains that can thrive in perchlorate-rich Martian soil without the necessity for perchlorate remediation (55).

The presence of perchlorates might be even beneficial for enzymatic activities at the low temperatures prevailing on Mars due to a reduced enthalpy of activation owing to chaotropic effects of perchlorate salts (67). Our data indicates that perchlorate-induced oxidative stress is not substantially higher than for other salts like NaCl. However, this might be only true for the Martian subsurface, because close to the surface, cosmic radiation could decompose perchlorates to far more reactive oxychlorine species such as hypochlorite which exhibit a strong oxidative stress (68) and be extremely detrimental to any life.

## Conclusions

The results of this study revealed perchlorate-specific microbial stress responses never described in this context before. Even though NaCl- and NaClO_4_-induced stress responses in *D. hansenii* share several metabolic features, we identified enhanced protein glycosylation, folding via calnexin cycle and cell wall biosynthesis as a counteractive measure to perchlorate-induced chaotropic stress which generally destabilizes biomacromolecules. At the same time, mitochondrial translation processes are downregulated under perchlorate-specific stress. When applying these physiological adaptations, cells can increase their perchlorate tolerance substantially compared to perchlorate shock exposure. These findings make it likely that putative microorganisms on Mars can draw on similar adaptation mechanisms enabling survival in perchlorate brines on Mars.

## Materials and Methods

### 5.1 Microbial cultures

The halotolerant yeast *D. hansenii* (DSM 3428) was obtained from the Leibniz Institute DSMZ - German Collection of Microorganisms and Cell Cultures. A stock culture was grown aerobically without shaking at 25°C (optimum growth temperature) in liquid DMSZ growth medium #90 (3% malt extract, 0.3% soya peptone) and was frequently reinoculated. Additionally, four different salt-containing liquid growth media (DSMZ #90) were prepared having a molal (m, mol/kg) salt concentration of either 1.5 m NaClO_4_, 2.4 m NaCl, 2.4 m NaClO_4_, or 3.9 m NaCl. The latter two concentrations represent the almost highest concentrations of the respective salt enabling growth of *D. hansenii*. The maximum growth-enabling concentrations reported to date are 2.5 m NaClO_4_and 4.0 m NaCl (13). We choose slightly lower concentrations to guarantee reproducible growth of the cultures and to generate sufficient biomass for protein extraction. The other two salt concentrations (1.5 m NaClO_4_and 2.4 m NaCl) represent moderate stress conditions of the respective salt (_∼_62% of the maximum salt concentration suitable for growth). The availability of two treatments with the same molal salt concentrations (2.4 m NaCl and 2.4 m NaClO_4_) allowed for an additional comparison of cellular stress responses to the two different salt species at the same osmolality. The growth media were prepared by mixing the media components, the salt and water, followed by pH adjustment (pH ∼ 5.6) and sterile filtration. All treatments (no salt, 1.5 m NaClO_4_, 2.4 m NaCl, 2.4 m NaClO_4_, and 3.9 m NaCl) were inoculated as biological triplicates, i.e. for each treatment three different samples were inoculated. The salt-free treatment as well as the samples containing 1.5 m NaClO_4_and 2.4 m NaCl were inoculated with the salt-free stock culture. Hence, the two saline treatments experienced a salt shock after inoculation. Since the respective salt shock would be too intense in 2.4 m NaClO_4_and 3.9 m NaCl treatments to enable growth, these samples were inoculated with long-term adapted cultures already grown at the respective salt concentration (Figure 1).

### 5.2 Sample preparation for proteomics

Protein extraction was conducted using the recently developed filter-aided Sample Preparation by Easy Extraction and Digestion (fa-SPEED) protocol (17). Cells were centrifuged for 3 minutes at 5.000 x g after reaching exponential growth phase which is 1 day for salt-free treatments, 3 days for 1.5 m NaClO_4_and 2.4 NaCl, 6 days for 2.4 m NaClO_4_, and 7 days for 3.9 m NaCl. Cell pelleting in 3.9 m NaCl samples was incomplete (turbid supernatant), but sufficient for further protein extraction. The reason for incomplete pelleting is presumably an electrostatic repulsion of cells because dilution of additional test samples did not result in larger pellets, but gently stirring with a grounded metal rod before centrifugation did. The cell pellets were washed three times with phosphate buffer saline (PBS) followed by cell lysing with 50 μL trifluoroacetic acid (TFA) for 3 minutes at 70°C. Afterwards, samples were neutralized with 500 μL 2 M tris(hydroxymethyl)aminomethane (TRIS) solution. After adding 55 μL reduction/alkylation buffer (100mM tris(2-carboxyethyl)phosphine / 400 mM 2-Chloracetamid), the samples were incubated at 95°C for 5 minutes.

Protein concentrations were determined by turbidity measurements at 360 nm using GENESYS™ 10S UV-Vis spectrophotometer (Thermo Fisher Scientific). Fifty μg proteins were diluted to 40 μL using a 10:1 (v/v) mixture of 2 M TrisBase and TFA, mixed with 160 μL acetone and incubated for 2 min at RT. For samples containing less than 50 μg proteins per 40 μL sample, the volumes of sample and acetone were increased at constant sample/acetone ratio until 50 μg protein/sample were reached. Afterwards, proteins were captured on Ultrafree®-MC (0.5 mL) centrifugal devices, 0.2 μm, PTFE (Merck) at 5000 x g for 2 min. The samples were washed successively with 200 μL 80% acetone, 200 μL 100% acetone and 200 μL n-pentane at 5000 x g for 2 min each.

Subsequently, 40 μL digestion buffer (50 mM ammonium bicarbonate) containing trypsin (1:25 (enzyme to protein ratio) Trypsin Gold, Mass Spectrometry Grade (Promega)) was added to the filter containing the proteins followed by incubation at 37 °C for 20 hours. The sample solution containing the digested proteins was centrifuged at 5.000 x g for 2 min and the filter was washed subsequently with 40 μL digestion buffer containing 0.1% TFA. 10% TFA solution was added until the pH of the samples reached approx. 2.Peptides were desalted using the Pierce™ Peptide Desalting Spin Columns (Thermo Scientific) according to the manufacture’s protocol no. 2162704. The desalted samples were dried in a vacuum concentrator. The dried peptides were dissolved in 0.1% formic acid and quantified by measuring the absorbance at 280 nm using an Implen NP80 spectrophotometer (Implen, Munich, Germany).

### 5.3 Liquid Chromatography and Mass Spectrometry

Peptides were analysed on an EASY-nanoLC 1200 (Thermo Fisher Scientific, Bremen, Germany) coupled online to a Q Exactive™ HF mass spectrometer (Thermo Fisher Scientific). One μg of peptides was separated on a PepSep column (15 cm length, 75 μm i.d., 1.9 μm C18 beads, PepSep, Denmark) using a stepped 30 min gradient of 80 % acetonitrile (solvent B) in 0.1 % formic acid (solvent A) at 300 nL/min flow rate: 5–11 % B in 2:49 min, 11–29 % B in 18:04 min, 29–33 % B in 3:03 min, 33–39 % B in 2:04 min, 39–95 % B in 0:10 min, 95 % B for 2:50 min, 95–0 % B in 0:10 min and 0 % B for 0:50 min. Column temperature was kept at 50°C using a butterfly heater (Phoenix S&T, Chester, PA, USA). The Q Exactive™ HF was operated in a data-independent (DIA) manner in the m/z range of 345–1,650. Full scan spectra were recorded with a resolution of 120,000 using an automatic gain control (AGC) target value of 3 × 10^6^ with a maximum injection time of 100 ms. The full scans were followed by 62 DIA scans of dynamic window widths using an overlap of 0.5 Th (16). DIA spectra were recorded at a resolution of 30,000 using an AGC target value of 3 × 10^6^ with a maximum injection time of 55 ms and a first fixed mass of 200 Th. Normalized collision energy (NCE) was set to 27 % and default charge state was set to 3. Peptides were ionized using electrospray with a stainless-steel emitter, I.D. 30 μm (PepSep, Denmark) at a spray voltage of 2.1 kV and a heated capillary temperature of 275°C.

### 5.4 Data Analysis and Statistical Information

Protein sequences of *Debaryomyces hansenii* (UP000000599, downloaded 16/10/20), were obtained from UniProt (69). A spectral library was predicted for all possible peptides with strict trypsin specificity (KR not P) in the m/z range of 350–1,150 with charge states of 2–4 and allowing up to one missed cleavage site using Prosit (70). Input files for library prediction were generated using EncyclopeDIA (Version 0.9.5) (71). The data was analyzed using the predicted library with fixed mass tolerances of 10 ppm for MS^1^ and 20 ppm for MS^2^ spectra using the “robust LC (high accuracy)” quantification strategy. The false discovery rate was set to 0.01 for precursor identifications and proteins were grouped according to their respective genes. The resulting pg_matrix.tsv file was used for further analysis in Perseus (version 1.6.5.0) (72).

The same program was used to z-normalize protein abundances followed by ANOVA (FDR = 0.01) and Post Hoc testing (FDR =0.05). Subsequently, the abundances of biological triplicates were median averaged, and the relative log2-fold changes of the salt-containing (saline) treatments compared to the salt-free control were calculated. The results were filtered for significant pairs of the salt-free samples and at least one of the saline treatments and were then plotted into a hierarchical clustered. Additionally, volcano plots have be generated with the same software after t-test of the z-normalized protein abundances. Protein groups of interest were annotated and analyzed with the STRING database (https://string-db.org/)(18) regarding enriched metabolic pathways and the formation of functional protein clusters.

## Acknowledgments and funding sources

This Research was funded by the Deutsche Forschungsgemeinschaft (DFG, German Research Foundation) – 455070607.

## Data availability

The mass spectrometry proteomics data have been deposited to the ProteomeXchange Consortium (http://proteomecentral.proteomexchange.org) via the PRIDE (73) partner repository with the dataset identifier PXD033237.

## Supplementary Information

The metabolic enrichments retrieved from the STRING database are summarized in Table S1. The general proteomic salt stress response of *D. hansenii* is shown in Fig. S1.

## Author Contributions

J.H. performed growth experiments with *D. hansenii*. J.H. and A.S. conducted protein extraction. J.H., J.D., D.M., and P.L. accomplished proteomic analysis. All authors, including J.H., J.D., D.M., A.S., P.L., H.-P.G., and D.S.-M., contributed to the interpretation of results. J.H. wrote the manuscript with input from all authors.

