## Supplemental Figure 1 for "Perchlorate-Specific Proteomic Stress Responses of *Debaryomyces hansenii* Could Enable Microbial Survival in Martian Brines"

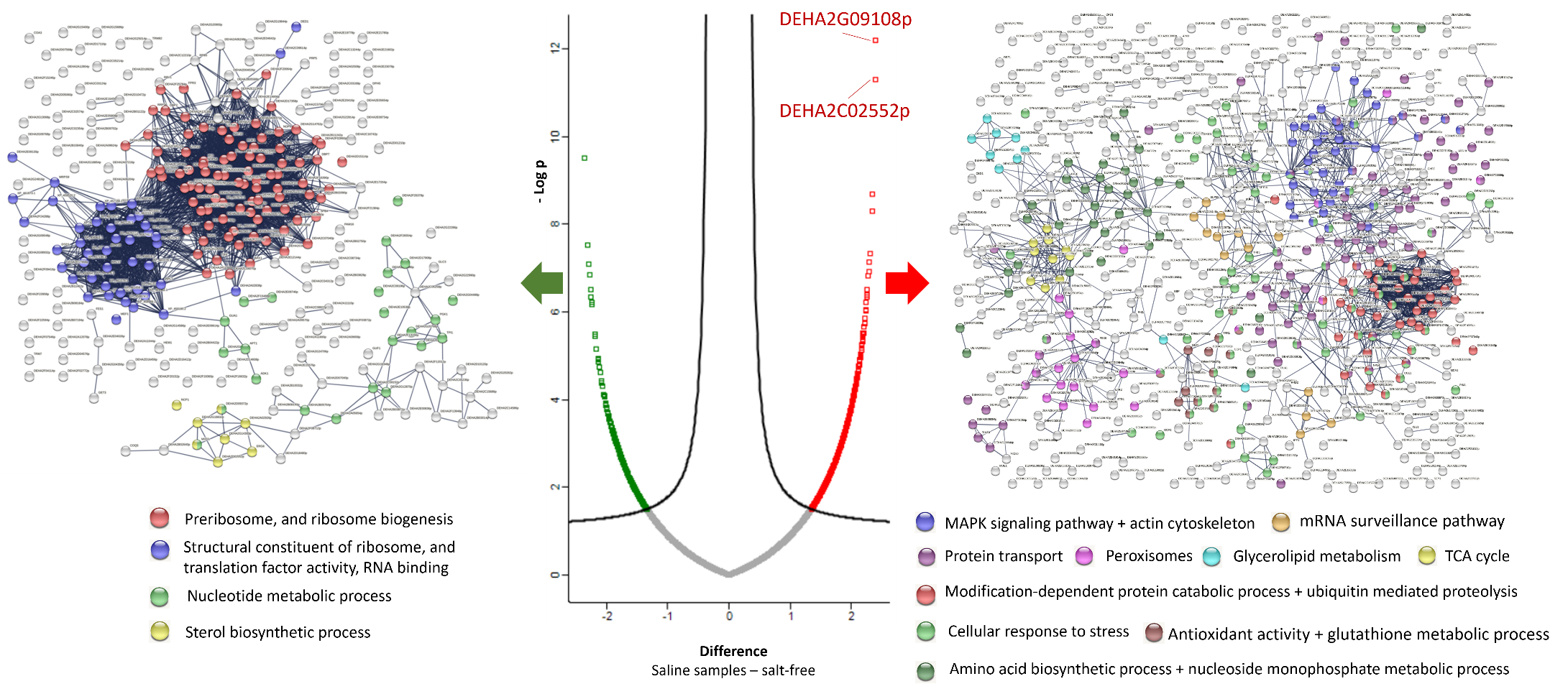

**Fig. S1:** General salt stress response. Volcano plot (at the center) of proteins significantly up- (red) or downregulated (green) in all salt-containing samples compared to the salt-free control, and STRING database analyses of up- (right) and downregulated (left) pathways. The most prominent protein clusters are color-coded, all other enrichments are summarized in Table S1. In the volcano plot the two ATPase-coupled cation transmembrane transporters DEHA2G09108p and DEHA2C02552p are labeled, which have the highest significance upon all upregulated proteins.
